# Item-specific clustering of high gamma-band activity on theta phase during speech categorization

**DOI:** 10.64898/2026.10.29.762424

**Authors:** Sanne Ten Oever, Joel Reithler, Mario E. Archila-Meléndez, Erik D. Gommer, Rob P.W. Rouhl, Vivianne H.J.M. van Kranen-Mastenbroek, Mark J. Roberts, Judith C. Peters

## Abstract

Theta-gamma phase-amplitude coupling (PAC) has been proposed as a mechanism for separating neural representations in time. Although item-specific PAC has been reported, its role in perception remains unclear. Notably, previous work has shown that theta phase can bias perception, raising the question whether such biases are linked to phase-specific gamma activity. Here, we investigated whether phase-dependent perceptual biases are accompanied by item-specific theta-gamma PAC. We presented ambiguous speech sounds that could be interpreted as either /da/ or /ga/ in patients with epilepsy implanted with intracranial electrocorticography. We found that pre-stimulus theta phase predicted both the neural and perceptual response, with /da/ and /ga/ tuned channels exhibiting gamma activity clustering at distinct theta phases. These findings suggest that neural and perceptual biases stem from phase-specific pre-activation of item-specific neural representations. They demonstrate the generalizability of the theta-gamma code, shaping behavior and enhancing representational capacity.

## Introduction

The phase of low frequency oscillations in the theta range has been shown to modulate the power of high frequency oscillations in the gamma range^1–3^. This theta-gamma phase-amplitude coupling (PAC) occurs in hippocampal^4,5^ as well as neocortical areas^1,3^, across species^3,6,7^ and tasks^8^. Theories propose that theta-gamma PAC organizes multi-item messages by representing different items at different oscillatory theta phases through phase-specific gamma power bursts^9,10^, resulting in a temporal separation which can disambiguate the representations. The organization of multiple representations has often been studied in the context of working memory where multiple items must be maintained simultaneously^4,5,11^. However, this multi-item theta-gamma PAC mechanism likely generalizes beyond working memory, as many neural functions rely on the communication and maintenance of multiple pieces of information ^9^.

Theta phase-dependent item-specific gamma is well established at the level of multi-unit and single-unit recordings^9,12,13^. Seminal work in rat hippocampus showed that the firing of place cells is locked to specific hippocampal theta phases that shift progressively as the animal traverses the place field, which changes as the experiment evolves, a phenomenon termed phase precession^14^. Phase precession has recently also been shown in single-neuron recordings in the human hippocampus^15,16^. Besides firing rates, gamma activity also clusters at specific theta phases^17,18^. This theta-phase clustering of gamma power bursts has been observed in human neocortex, for example during picture viewing^17^ and spatial navigation^18^. These studies reveal item-specific coupling, but do not establish its functional relevance for behavior.

The functional relevance of theta-gamma PAC for multi-item processing is supported by covariation between theta-gamma PAC strength and behavioral performance fluctuations. In rat hippocampus, stronger PAC leads to improved performance in both spatial memory and item-content associations tasks^19,20^. In humans, PAC-behavior relationships have been reported across a variety of tasks including temporal order^21^, association^22^, working memory^5,23,24^ and long-term memory ^25^. Additionally, transcranial alternating current stimulation in frontal cortex at theta-gamma frequencies improves memory performance, suggesting a causal role^26^. Together, these studies demonstrate the behavioral relevance of theta-gamma PAC in memory tasks.

As the brain cycles through the different neural messages to maintain information, new inputs may be processed differently depending on the theta phase at which they arrive. Direct behavioral evidence for such phase-dependent processing comes from electroencephalography/ magnetoencephalography (EEG/MEG) studies showing that the phase at which an ambiguous stimulus is presented biases perception toward one of two interpretations^27–31^. This effect is hypothesized to arise from theta-gamma PAC, in which gamma activity for the two perceptual representations occurs at distinct theta phases ^32^. Because ambiguous input activates both possible representations, the representation that is active at the moment of input arrival (i.e. at that phase) has a slight advantage and is activated more strongly, biasing behavior toward that representation. While a theta phase-dependent bias has been demonstrated^27^, the association with gamma bursts is unknown as non-invasive methods cannot measure gamma activity in the high frequency ranges. Monitoring gamma power bursts can only be recorded close to its neural source, which requires intracranial recordings. Thus, the hypothesis that this phase-dependent perception bias is accompanied by item-specific clustering of gamma activity at different theta phases^32^, remains untested.

Here, we investigated whether phase-dependent perceptual bias is accompanied by item-specific theta-gamma PAC. We ran a speech categorization experiment^27^ in eight patients with intractable epilepsy, undergoing continuous electrocorticography (ECoG) recording for pre-surgical evaluation. Ambiguous sounds that can be perceived as either /da/ or /ga/ were presented along with non-ambiguous /da/ and /ga/ sounds. In stimulus-responsive channels, the phase of ongoing theta oscillations at sound onset biased the perception of ambiguous sound toward /da/ or /ga/^27,29^. Likewise, this pre-stimulus theta phase biased the neural activity profile of the ambiguous sound toward a /da/ or /ga/ representation. Next to these phase-dependent biases, we observed that the theta phase at which high-gamma activity clustered was related to the tuning of individual recording sites for /da/ or /ga/. These findings suggest that theta-gamma PAC temporally segregates competing neural representations. Additionally, they indicate that this mechanism extends beyond memory to perception, thereby underscoring the generality of theta-gamma PAC.

## Results

### Ambiguous and unambiguous sound perception

We used previously generated morphs between /da/ and /ga/^27^. They were created by incrementally transforming a recorded auditory /da/ in a /ga/ through shifts of the third formant from the original frequency band of 3kHz down to 2.6 kHz (in nine incremental steps). Per session, we presented 180 stimuli per session pseudo-randomly (60 extremes and 120 ambiguous sounds) and participants had to indicate which sound they heard. We extracted the behavioral profile by calculating the proportion /da/ responses to all sounds. Overall, our selection of participants could differentiate both between the extreme /da/ and /ga/ sounds, as well as between the extreme sounds and the ambiguous morph (Figure 1A and S1; Kruskal-Wallis H(2,21) = 17.37, p < 0.001; all follow-up sign rank tests: Z = 36, p = 0.008), indicating they could do the task.

**Figure 1.**
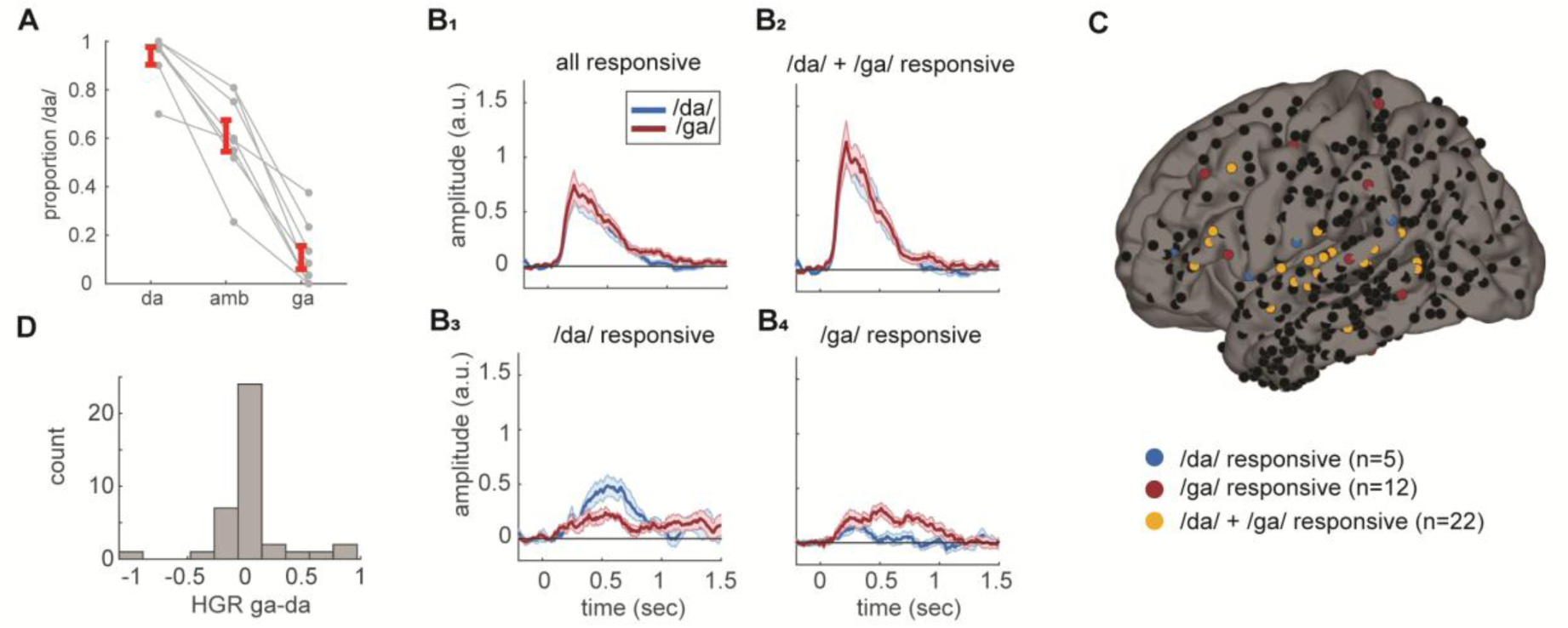
Behavioral results and responsive channels. A) proportion /da/ responses for the /da/, /ga/, and ambiguous (amb) sound. Gray lines represent individual participants; red lines represent the group mean with its standard error (Kruskal-Wallis H(2,21) = 17.37, p < 0.001). B) High gamma band activity for /da/- responsive, /ga/-responsive, and /da/ + /ga/ responsive channels. Shaded areas represent the standard error of the mean. C) Location of responsive channels plotted on an average brain. D) Number of responsive channels as a function of high-gamma-response (HGR) difference between /da/ and /ga/ in an early stimulus evoked window (0.05-0.25 s).

### Stimulus-responsive channels in lateral cortex

First, we aimed to identify channels that responded to our sounds. We extracted trial-by-trial high gamma responses (HGR; 80-160 Hz; see methods for details) and compared post-stimulus activity from a 0-1000 ms window to baseline (average of -500 – 0 ms window) using a sign rank test and correcting for multiple comparison across time points using cluster-based statistics^33^. A total of 39 channels responded significantly to one or both sounds across six of our recorded patients (total of 463 channels; Figure 1B-D). Note that the channels that were identified to only respond to one sound seemed to also have a (non-significant) response to the other sound (Figure 1B). However, the responses of these channels were overall lower in amplitude than those in channels that significantly responded to both sounds. The non-selectivity of our channels is not surprising considering that the sounds are very similar and only differ in the frequency of the third formant. The responsive channels were mostly located in the left superior temporal cortex (n=12) but were also present in left supramarginal (n=4) and left pars opercularis (n=4; see Supplementary Table 1 for a full overview). Next to these 39 channels, we identified the 39 least responsive channels used for control analyses (Figure S2 and methods).

### Phase-dependent perception of ambiguous sounds

Next, we identified whether the phase of ongoing low-frequency oscillations in the responsive channels was linked to the behavioral response, such that participants were more likely to perceive /da/ or /ga/ dependent on the response, similarly as we previously reported (see ^27^ and ^29^). We extracted low-frequency phase (1-15 Hz) by performing wavelet analysis over a time window from -1 up to 0 second around stimulus onset for all ambiguous sounds. Critically, for the time-frequency analysis we extracted the phase of ongoing oscillations by padding the post-stimulus time interval of each trial with the value of its last timepoint, ensuring that activity from the post-stimulus window did not smear into the pre-stimulus period, while avoiding edges and high frequency artifacts. To test for a phase-response relation, we performed a logistic regression for each channel with the sin and cosine of the ongoing phase per trial as dependent variable and behavioral choice (/da/ or /ga/) as independent variable. We obtained the χ^2^ statistic from the likelihood ratio test by comparing this model with an intercept model (Figure S3). We then generated the expected χ^2^ distribution by shuffling the behavioral choice labels and repeated the logistic regression and null-model comparison across 1000 permutations. Subsequently, we transformed the skewed χ^2^ using a boxcox transform (henceforward called χ^2^*) and used the mean of the null distribution as the expected value. Expected and observed values were entered in a second level multi-level linear mixed effect model (lme) with χ^2^*-type (observed vs expected) as fixed factor and channel and participant random factors. We compared this model to an intercept model and corrected post-hoc multiple comparisons using a cluster-based permutation approach, in which the χ^2^*-type labels were randomly permuted^33^.

We identified a significant cluster (p=0.013) in which pre-stimulus phase predicted participants’ response (Figure 2A). This cluster had a frequency range between 1.95 and 6.47 Hz and a time range of -0.81 and -0.55 s before stimulus onset. These effects are similar to those we previously reported, although at slightly lower center frequency^27,29^. Note that such effect might be expected near time zero, but detecting it is challenging because the data were padded after stimulus onset. Since wavelet estimation requires a temporal window, estimates zero inevitably include contributions from the padded segment (see dotted line in Figure 2). We found no significant effect for the 39 least responsive channels (lowest cluster p-value=0.830). These findings suggest that the oscillatory phase in sound-selective brain areas determined how participants interpreted the ambiguous stimulus.

**Figure 2.**
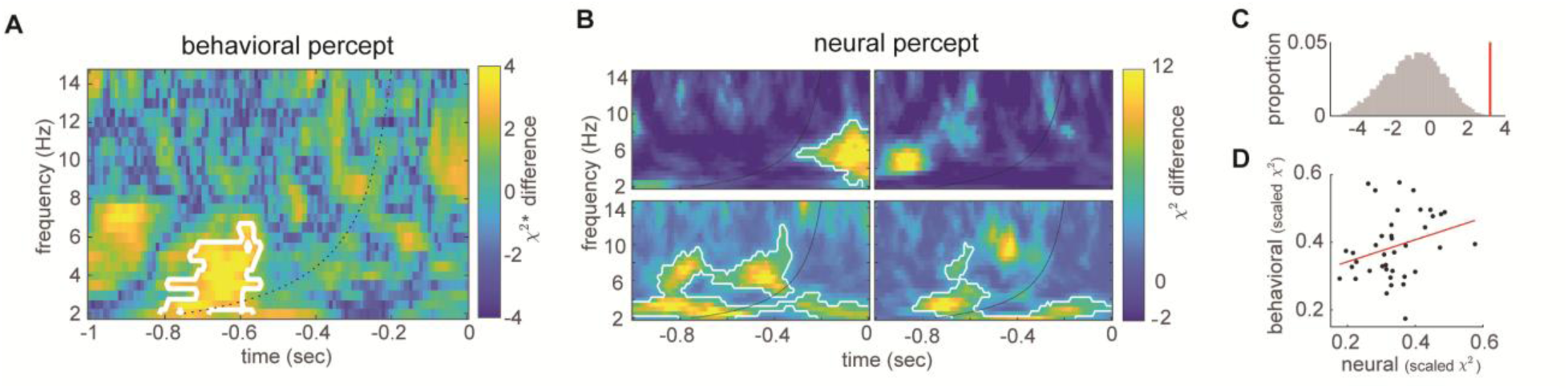
Pre-stimulus phase determines the behavioral and neural percept of an ambiguous sound. A) Behavioral percept. The outcome of the observed minus the expected χ^2^* from the lme model is plotted across frequency and pre-stimulus time for the responsive channels. Results show that oscillatory phase determines the behavioral percept (/da/ or /ga/). White outlines indicate borders of significant clusters (p=0.013). All data left of the black dotted line have no padded data (post-zero) included in the estimate. B) Neural percept. Outcome of the observed minus the expected χ^2^ from the logistic regression for example channels. White outlines indicate clusters significant at the single channel level (p < 0.05). The layout is identical to A. C) Permutation test of the effect of pre-stimulus theta phase on the neural percept across channels, within the significant cluster of the behavioral percept analysis. Gray bars show the proportions of permutations per bin of the null distribution of the second-level χ²* statistic (10,000 permutations). The red line marks the observed statistic (χ²* = 4.053; p = 0.0007). D) Relationship between behavioral and neural percept effects across channels (Spearman’s ρ = 0.32; p = 0.0482). Each dot represents the channel’s mean min-max normalized first-level χ² values of pre-stimulus theta phase, within the significant cluster of the behavioral percept analysis (Figure 2A), for the behavioral and neural percept. The red line shows a linear fit to the dots.

### Phase-dependent neural representations of ambiguous sounds

Given that pre-stimulus phase influences the behavioral, we next asked whether it also shapes the neural response to ambiguous sounds. To this end, we trained channel-specific linear support vector machines (SVMs) to discriminate HGRs elicited by the extreme /da/ and /ga/ sounds and used these classifiers to predict the *neural percept* (i.e., the predicted class label) of each ambiguous sound. We tested the dependence of this neural percept on pre-stimulus phase by using analyses analogous to those in the previous section, with the neural percept replacing the behavioral percept as the dependent variable in the first-level single-channel logistic regression. These results showed significant single-channel effects of pre-stimulus theta phase, with a diversity in exact timing of the effect across channels (Figure 2B; see channels split by brain region in S4).

Across-channel significance was investigated specifically for the cluster of the behavioral percept to assess whether pre-stimulus phase influences neural and behavioral percepts similarly. We applied the second-level lme model to the time–frequency points within the significant cluster identified in the behavioral percept analysis (Figure 2A). Pre-stimulus theta phase strongly influenced neural perception in this cluster (χ^2^* = 4.053; p = 0.0007; 10,000 permutations; Figure 2C). Moreover, across channels, the first-level χ^2^values within this cluster were positively correlated between the behavioral and neural percept analyses (Spearman’s ρ = 0.32; p = 0.0482; Figure 2D).

### Theta-gamma phase-amplitude coupling

To verify the presence of theta in our data, we extracted the power spectrum across all responsive channels for the -1 to 0s pre-stimulus window (Figure 3A). We could identify a peak around 6.84 Hz. This frequency has more often been reported in middle temporal cortex, where most of our responsive channels are located^29,34,35^. Then we calculated phase-amplitude coupling in the same window using the modulation index^36^ across 1-15 Hz low-frequency phase and 30-160 Hz high-frequency power (for details see methods). For statistical analysis we again performed permutations to extract the expected and observed modulation index values, as described above in the phase-dependent perception of ambiguous sounds. Again, we performed a second level analysis on these expected and observed modulation index using lme in combination with cluster-based correction. Additionally, we conducted single channel statistics using the modulation index permutations for cluster-based correction on the single channel level.

**Figure 3.**
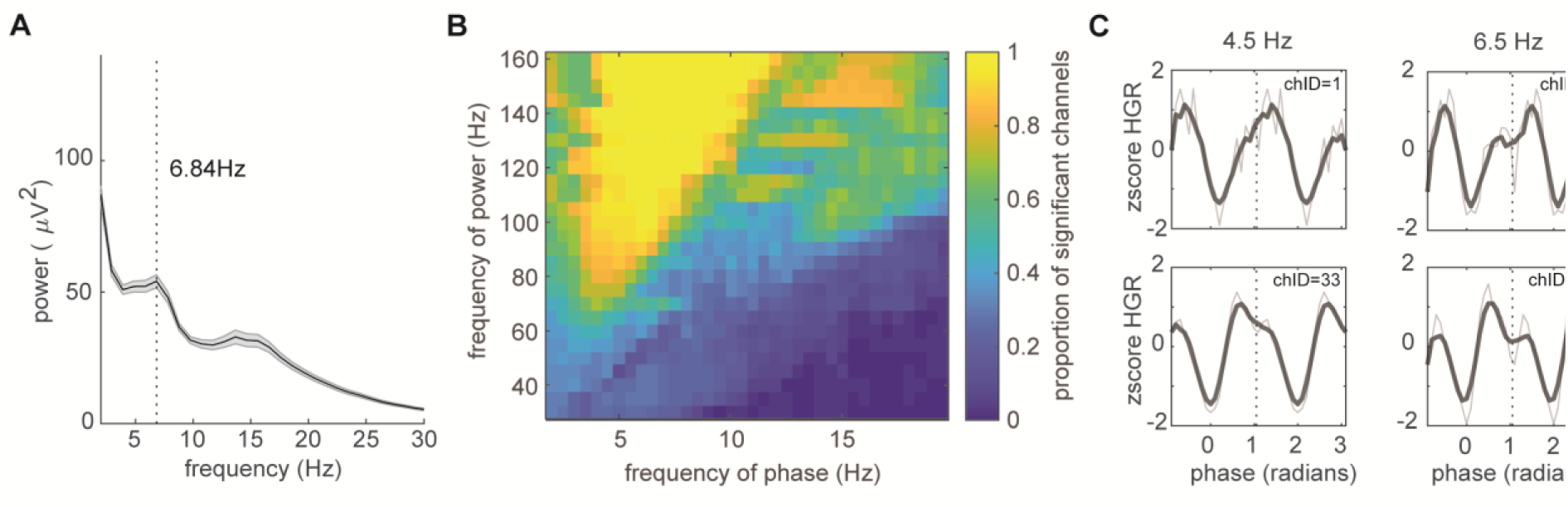
Phase-amplitude coupling. A) Power spectrum in the -1 to 0 pre-stimulus window for the responsive channels. Error bar indicates the standard error of the mean across channels. B) Phase-amplitude coupling results for the single channel analysis. Proportion of channels with a significant cluster are visualized. C) Average z-scored high gamma response (HGR; 80-160 Hz) across different phase bins for 4.5 and 6.5 Hz for two representative channels. Gray lines represent the raw traces, and black lines indicate the 5-point moving average.

We found significant phase-amplitude coupling across almost all channels (Figure 3B), with the largest proportion of channels having a cluster between theta (mean highest MI* across channels at 7.4 Hz; standard deviation of 4.25 Hz) and high gamma (mean highest MI* across channels at 144.14 Hz; standard deviation of 34.43 Hz). On the second level analysis, this cluster encompassed large range of the phase and power points, also with the strongest effect in the theta-high gamma range (Figure S5). These data suggest that indeed, there is strong theta-gamma phase-amplitude coupling. We next focused specifically on this theta-gamma coupling by only taking the phases of the theta range (2-8 Hz) and the average of the power spectrum for all frequencies between 80 and 160 Hz. Figure 3C shows examples of phase-amplitude coupling in the high gamma range at 4.5 and 6.5 Hz for two different electrodes. What is interesting is that the peak of the phase-amplitude coupling is not always at the same phase. The next analysis aims to investigate whether this systematically relates to the tuning of the channel to either /da/ or /ga/.

#### /da/-/ga/ tuning relates to the theta phase at which gamma clusters

In the previous analysis we saw that gamma clustered at different phases of theta for different channels. To investigate whether this relates to the tuning of the channel to either /da/ or /ga/, we related the phase at which pre-stimulus gamma had the strongest value (across the 20 phase bins) to the average difference of the high-gamma response between /da/ and /ga/ for the early sensory response period (0.05-0.25 s). We plotted all data in a circular plot, with as radius the order of the absolute difference, and the color the signed difference (Figure 4A). A picture arises in which for 6.5 Hz, /da/ sensitive channels cluster at -0.5 π, while /ga/ sensitive channels cluster around 0.5 π. For the statistical test, we performed a lme with sin and cosine of the theta peak phase as dependent variables and gamma response difference as independent variable. This was repeated for all theta frequencies in the range of 2-8 Hz. We found significant effects across the theta range peaking at 3.0, 4.5 and 6.5 Hz (FDR corrected; Figure 4C; see Figure S6 for the rose plot for 4.5 Hz; at 6.5 Hz χ²=14.27; FDR-corrected p = 0.0087). A similar pattern of results remained when using the common averages instead of the average of the electrodes-strip as offline reference.

**Figure 4.**
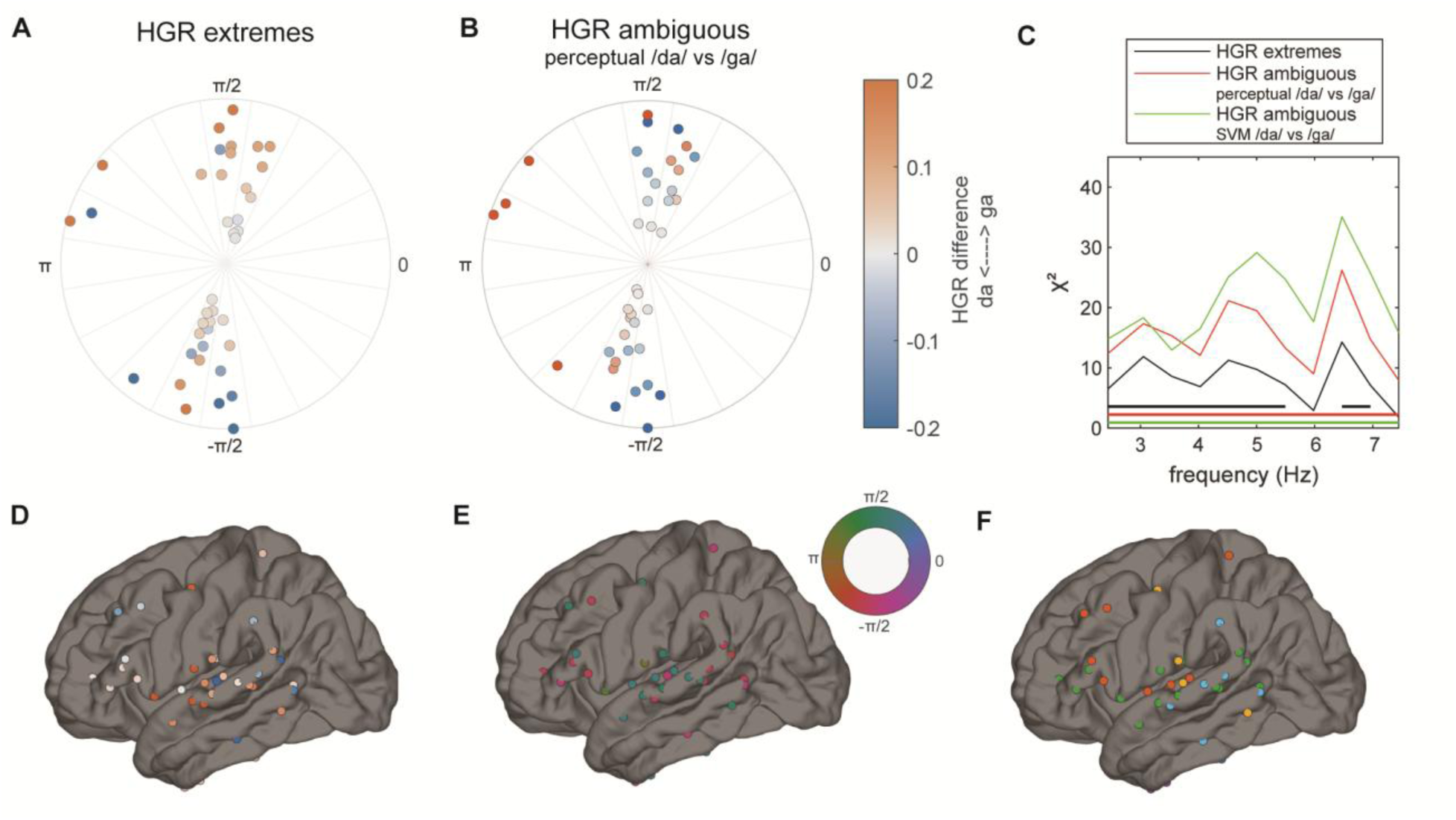
Channel tuning relates to gamma clustering phase. A) Rose plot displaying the peak gamma phase for each channel together with the tuning of that channel (orange indicates /ga/ and blue indicates /da/). Tuning is defined the difference between the extreme /da/ and /ga/ sounds. Radius represents the absolute difference between /da/ and /ga/ responses. B) Same as A but for the difference in perceived /da/ and perceived /ga/ for the ambiguous sounds. C) LME results across all tested theta frequencies for tuning to extreme sounds, the ambiguous sounds, or the SVM classification of ambiguous sounds. Thick lines indicate significant effects (p < 0.05) for the corresponding color-coded contrasts. D) Cortical distribution of the phoneme tuning (/da/ vs /ga/) of the channels, color scheme is the same as in A and B. E) Distribution of the peak gamma clustering phase. F) Distribution across participants. Different colors indicate different participants.

After observing this relation between gamma clustering phase and tuning to extreme sounds, we asked whether a similar association exists between gamma clustering phase and tuning to the perceptual interpretation of ambiguous sounds. To this end, we repeated the previous analysis, but now with channel tuning strength defined as the HGR difference between ambiguous sounds perceived by the participant as /da/ versus /ga/ or classified as /da/ versus /ga/ with the SVM. A similar profile appeared as for the relation between phase clustering and tuning to extreme sounds with significant effects found across the theta range peaking at 3.0, 4.5 and 6.5 Hz. This was irrespective of whether the tuning was defined by the behavioral percept or by the SVM prediction of the ambiguous sound (Figure 4B,4C): At 6.5 Hz, clustering was significant for tuning defined by the behavioral percept (χ²=26.22; FDR-corrected p = 0.0002) and the SVM prediction (χ²=35.09; FDR-corrected p = 0.00002; Figure S6).

To investigate the spatial distribution of this effect, we made topographical plots color coded for phoneme tuning to extreme sound per channel (Figure 4C), peak gamma phase (Figure 4D), and patient ID-number (Figure 4E). None of these topographies showed coherent spatial clusters of tuning or a mapping of gamma clustering peak phase.

## Discussion

In the current study we investigated whether high-gamma activity clusters at distinct theta phases during a speech categorization task known to induce theta phase-dependent behavioral biases^27^. We presented ambiguous auditory stimuli that could be interpreted as a /da/ or a /ga/ to patients with epilepsy implanted with ECoG and asked them to identify the sounds. We found that pre-stimulus theta phase biases both the participants’ perception of ambiguous sounds as /da/ or /ga/ and the corresponding neural percept decoded from high-gamma responses. Additionally, gamma phase clustered at different theta phases for channels tuned to either /da/ or /ga/, respectively. Interestingly, this held true for tuning in both the acoustic domain (extreme sounds) as well as the perceptual domain as reflected by the behavioral and neural classification of ambiguous sounds. These results show that previously reported phase-dependent biases go together with item-specific theta-gamma PAC both for tuning in the acoustic and perceptual domain, suggesting that the bias is a result of pre-activating representations at distinct oscillatory theta phases. This highlights the importance of theta-gamma PAC for behavior.

Although the presence of item-specific theta-gamma PAC has been demonstrated before^11,14,17,18^, it has not yet been linked to phase-dependent behavioral biases. This link reveals an important consequence of time-resolved item representations: phase separation can expand representational space, but it does cause a phase-dependent bias when information is ambiguous. Our previous work showed that stronger categorical perception relates to increased phase-separation^28^, suggesting that phase-coded representations benefit behavior. Conversely, a strong phase-dependent bias in perception could be dysfunctional, because phase rather than stimulus information would then determine the percept. Such biases could nonetheless be functional when stimulus content depends systematically on timing^37^. For example, more predictable words tend to be uttered earlier ^32,38,39^. This temporal dependency could be used to sort speech content on specific theta phases, so that each item is represented at the phase at which it is expected to arrive^32^.

It remains an open question what principles govern the clustering of gamma activity at different oscillatory phases. Most existing theories propose mechanisms based on the temporal order of events^16,40^ or their spatial position^14,41,42^. However, in the current paradigm, there is neither a temporal nor spatial structure to the input. The only difference between the two stimuli lies in their spectral content and their natural statistical distribution in the environment (/da/ is more frequent than /ga/). Phase-dependent biases have been linked to differences in statistical likelihood^29^, suggesting that gamma clustering may similarly reflect these likelihood differences. Such an ordering principle could arise as stimuli that are more frequent have a higher excitable representation, making them more likely to be active even at less excitable phases^43^. Whether this excitability-phase relationship extends to phase-dependent codes observed in temporally or spatially structured input or is specific to the perceptual cortical codes reported here remains an open question.

Original reports of item-specific phase-amplitude coupling have identified strong effects in hippocampus^14,15^. This is expected considering that the tasks used in these studies target hippocampal activation, for example spatial navigation or associative learning. Moreover, many of these studies only measured in hippocampus. The presence of theta-gamma PAC in cortex has been established^1^. But only a few studies explored item-specific PAC^11,17,44^ and those studies mostly targeted the mechanisms of working memory instead of behavioral biases. In our study, the hippocampal electrodes did not strongly respond to the stimuli, thus we could not investigate phase-dependency in hippocampus with the current task design. Nonetheless, we find item-specific theta-gamma PAC in regions associated with auditory sensory processes, such as superior temporal cortex. The frequency of the effect is comparable to cortical item-specific theta-gamma clustering^11,17^, and consistent with the reported theta frequencies in auditory regions^3,29,45^. Note that while elevated phase-amplitude coupling has been reported in patients with epilepsy^46^, the item-specificity of this effect is unlikely to be related to the pathology. Instead, these results highlight the generalizability of theta-gamma PAC as a neural code beyond memory-related tasks.

Research on phase-dependent behavioral responses has mostly focused on the idea that phase modulates excitability and consequently modulates overall performance (see e.g. ^47^). However, this reasoning passes by the observation that different neural populations can be active at distinct neural phases. If so, one should expect content-specific phase-dependent effects like we report here (also see ^48^). Recently, phase-dependent effects have been observed across different brain locations linked to spatially moving oscillatory patterns, called travelling waves^49–51^. However, those results are still interpreted under the assumption that optimal processing necessarily happens at the most excitable phase; this phase just happens to move across cortical locations representing different locations^52^. Future research should investigate how item-specific clustering affects item-specific behavioral processes.

Based on our significant electrodes one might believe that our effect is lateralized to the left hemisphere. We believe this interpretation requires caution as our sample was small. Within this sample, we had three patients with extended left cortical coverage and two with extended right cortical coverage (the other patients had bilateral temporal strips, mostly covering the middle and inferior temporal gyrus as well as depth electrodes). One of the right-lateralized implants (third row in figure S7) is also located rather posterior, covering more parietal-occipital than temporal cortex. In this patient, we only find the most anterior electrodes to be responsive, suggesting that if the implant was more anterior, we might have found more responsive channels. However, for one patient with a right lateralized implant that did cover the biggest portion of STG (where most significant electrodes were found [see Supplementary Table 1]) no responsive channels were found. Previous reports do suggest a lateralization of language processing to the left hemisphere^53,54^. This lateralization mostly relates to higher order lexical and syntactical processes, rather than low-level syllabic processes that we target here. Indeed, in a previous study using similar stimuli, we did not find strong evidence for lateralization^29^. However, task demands and cognitive task could still trigger left-lateralization even with the use of low-level phonetic stimuli^55,56^. With the current sample size, it is an open question whether our task involves lateralization.

The brain needs to represent multiple neuronal messages at the same time. Item-specific theta-gamma PAC is a candidate mechanism to separate these messages avoiding interference^9^. Here, we demonstrate that item-specific theta-gamma PAC is present in a perceptual task, primarily in superior temporal cortex. This clustering is accompanied by phase-dependent behavioral biases such that the time or phase of presentation determines the percept of the participant. This suggest that even in simple perceptual tasks, there is clustering of neural representations, something previously mostly associated with more complex cognitive tasks such as working memory^9,11^. This study highlights the importance and generalizability of item-specific theta-gamma PAC across the brain but also shows that this item-specific clustering mechanism can cause a bias in behavioral responses.

## Methods

### Participants and intracranial recordings

We included eight participants (5 females, 3 males, average age 39,5 years) with refractory epilepsy, who were monitored at the Epilepsy Monitoring Unit of the Maastricht University Medical Center. All participants were native Dutch speakers and could reliably identify the auditory stimuli (we tested three additional participants, but they were unable to do the task). Patients underwent continuous ECoG recording with standard subdural electrodes (Ad-Tech Medical Instrument Corporation, Oak Creek, WI, USA), as well as (hippocampal) depth electrodes recordings as part of their pre-surgical evaluation for possible resective epilepsy surgery (total channels: 580; range: 48-90). The electrode locations were based entirely on clinical criteria, consistent with prior reports ^57^. Data were recorded using BrainRT software (version 2.0.3164.0, OSG BVBA, Rumst, Belgium) at a sampling rate of either 1024 or 2048 Hz with mostly a superficial frontal EEG electrode as an online reference which showed limited epileptic activity during seizures. Data was amplified using Brainbox EEG-1166 amplifiers (Braintronics B.V., Almere, Netherlands). Participants were invited to join the study during preoperative consultation and were given no financial incentive to participate. All gave written informed consent before inclusion into the study. The task was performed during the first five days after surgical implantation, clinical condition and patient situation allowing. The study complies with the Declaration of Helsinki and was approved by the Medical Ethical Committee of the Maastricht University Medical Center, Maastricht, The Netherlands (METC number 14-2-004).

### Stimuli

The original stimuli were created by incrementally transforming a recorded auditory /da/ in a /ga/ sound by shifting the third formant from the original frequency band of 3kHz down to 2.6 kHz in steps of -19 Mel in Praat^58^, resulting in nine stimuli (see ^27^ for exact procedure). In the current study we used morph numbers 1 and 9 to represent the extreme sounds and morphs 3, 4, 5 for the ambiguous sounds (for the first two participants we set the morphs to 4, 5, and 6). These sounds were selected based on prior findings that most participants showed ambiguous morph boundaries near these values, thereby obviating an additional session to individualize these sounds (see Figure 1A). Sounds were presented using Presentation software (Neurobs) via the speaker of the laptop at a comfortable volume for the participant.

### Procedure

First, participants were familiarized with the stimuli by repeatedly presenting the extremes (clear /da/ and /ga/) and having participants indicate whether they could identify them. If participants were able to identify the extremes they could proceed to the main experiment. In the main experiment we presented the five stimulus types (two extremes and three levels of the ambiguous sounds) in random order and asked participants to identify the sounds by pressing corresponding buttons. The intertrial interval varied between 1.5s and 2.5s (relative to the response onset). For each session there were 180 stimuli, 60 extremes, and 120 ambiguous sounds (30 per level of the extremes and 40 per level of the ambiguous sounds). After every 60 stimuli, participants were encouraged to take a break. Five participants completed 1 session, two completed 2 sessions, and one completed 3 sessions.

### Anatomical segmentation and Intracranial electrode localization

Presurgical T1-weighted MRI scans were initially segmented using FreeSurfer (version 6.0)^59,60^. Then, quality control was performed through visual inspection. The remaining errors were manually corrected using ITK-SNAP software (version 3.4.02; ^61^. The postsurgical CT scan was co-registered to the presurgical T1-weighted MRI using Advanced Normalization Tools^62^. We used the Yael toolbox to spatially localize the intracranial electrodes in the MRI space^63^. Subdural electrodes were projected to the nearest point on the dural surface reconstructed from the presurgical MRI to correct for post-implantation brain shifts^64^. Finally, for visualization purposes, electrodes of all patients were projected to the brain average surface of FreeSurfer.

### Data analysis

#### Preprocessing

All analyses were performed in MATLAB (Mathworks) using Fieldtrip (version 20180712 for preprocessing, and version 20220707 for all other steps; ^65^) the circular toolbox^66^, and custom-made MATLAB scripts. First data was demeaned, bandpass filtered from 0.2-200 Hz and we applied a notch filter at 50, 100, and 150 Hz. Channels in the seizure onset zone were removed (mean = 5.33; range: 0-20) and additional bad channels were removed based on visually inspection (mean = 2; range: 0-13). Data was re-referenced to the average of the grid, strip, or stereotactic lead of which each channel belonged. Then data was cut from -2 to 3 s around sound onset and downsampled to 500 Hz. Trials deviating more than 3.5 standard deviations from the mean, or exceeding 300 μV, were excluded.

#### Time frequency analysis

Time-frequency was calculated separately for the pre- and post-stimulus analysis. For both analysis we extracted low-frequency Fourier spectra for a frequency window of 2 to 30 Hz (in steps of 0.5 Hz) using a wavelet approach with a linear spacing of 3 to 6 cycles across the frequencies of interest. For the high frequency analysis, we used a multi-taper convolution approach using discrete prolate spheroidal sequences across frequencies from 30 to 160 Hz, in 5 Hz time steps. The time window length was set to include 10 cycles per frequency, with spectral smoothing set to half the target frequency. For the post-stimulus analysis, the time window of interest varied between -0.2 to 1.5 s with a temporal resolution of 10 ms. For the pre-stimulus analysis, the time window of interest was -1 to 0 s with a 10 ms resolution. To ensure that post-stimulus data did not influence the pre-stimulus window, we padded the data beyond time zero with the value at stimulus onset for this analysis.

#### Evoked high gamma responses (HGR)

First, we identified the channels that were responsive to the extreme /da/ and /ga/ stimuli. To this end, we extracted all trials where participants correctly identified the /da/ sound. We subsequently used one-tailed sign rank Wilcoxon tests to investigate if the post-stimulus high gamma response (80-160 Hz; 0-1 s) was significantly stronger than baseline (average between -0.5-0 s). This test was corrected for multiple comparisons across time using cluster statistics^33^. Across channels we corrected for multiple comparisons using False Discovery Rate corrections (with a proportion of false discoveries of q > 0.05) for the strongest cluster per channel. We repeated this procedure for the trials with correctly identified /ga/ sounds. Only sound responsive channels (i.e., those responding to at least one stimulus) were included in the main analyses. For comparison, we selected an equal number of non-responsive channels, defined as those with the lowest absolute mean difference from baseline (0-1 s post-stimulus).

#### Phase-dependent categorization of behavioral percepts

For the sound-responsive channels we investigated whether phase influenced whether ambiguous stimuli were perceived as one or the other sound. We took the pre-stimulus Fourier spectra, extracted the phase, and performed per channel logistic regression (binomial distribution) using the sine and cosine of pre-stimulus phase as the independent, and response label (/da/ or /ga/) as dependent variable^67^. We selected time windows ranging between -1 and 0 s and frequencies of 1 – 15 Hz. This model was compared to an intercept model, and we extracted the observed χ^2^ of this comparison. We then created a null distribution per channel by permuting the response labels across trials enabling us to extract expected χ^2^ values (n = 1000). As the χ^2^ is skewed to the right, we performed a boxcox transform (the lambda was estimated from the permuted χ^2^ values), thereby normalizing all χ^2^ values (now called χ^2^*). The expected χ^2^*-value was defined as the mean of the null distribution. Then we performed a multi-level linear mixed effect (lme) analysis using the intercept of the channel, participant, and their interaction as random factors. We chose a lme model to ensure that we could model the dependency of channel and participant as some participants have multiple channels in the analysis (included in the random effects; also see^68^). As a fixed effect, we used the χ^2^*-type (observed or expected). This was compared to a model with the same random factors, but only the intercept as a fixed factor, again extracting χ^2^ with a likelihood test. This was repeated for all time-frequency points. To correct for multiple comparisons across time-frequency points, we performed cluster-based correction ^33^. To do so we repeated the multi-level linear mixed model using permutations for the χ^2^-type factor to generate a null-distribution needed for the cluster-based statistics (n=1000). Again, we corrected the χ^2^ resulting from the comparison to the null-model with a boxcox transform. The sum of the χ^2^* was used as our dependent variable with an alpha set to 0.05 (see Figure S3 for overview of the analysis steps).

#### Phase-dependent categorization of neural representations

To examine whether pre-stimulus phase influenced neural representations of ambiguous stimuli, we decoded /da/ and /ga/ percepts from HGRs in sound-responsive channels. We then tested the dependence of these percepts on pre-stimulus phase using analyses analogous to those in the previous section (Figure S3).

For each channel and trial, we averaged power in each of the 16 high gamma bands (80–160 Hz) over 0–250 ms post-stimulus onset, separately for extreme and ambiguous sounds. This yielded two trial × band matrices per channel: one for extreme /da/ and /ga/ sounds (extreme HGRs), used to train *extreme HGR SVMs*, and one for ambiguous sounds, split by the reported percept (ambiguous HGRs), used to train *ambiguous HGR SVMs*. Both SVM types were trained with the same procedure, using linear soft-margin SVMs per channel (*fitcsvm* function in MATLAB Statistics and Machine Learning Toolbox, v2026a), with L2-regularized weights and hinge loss, which penalizes margin violations in proportion to the box constraint C (controlling how strongly a model penalizes classification errors). To avoid overfitting through hyperparameter tuning on small samples and ensure consistency across training repetitions, C was fixed at its default (C=1). To prevent bias toward the majority class, we equalized /da/ and /ga/ trial counts in the training set: the majority class was randomly partitioned into non-overlapping subsets the size of the minority class, and any incomplete final subset was filled with trials randomly drawn, without replacement, from the other subsets. An SVM was trained on each subset together with the minority-class trials, so that every majority-class trial entered at least one training set. This partitioning was repeated five times with different randomizations. Within each training set, features were z-scored with that set’s mean and standard deviation, which were also applied to the test data. Extreme HGR SVMs were trained on all extreme HGR trials and applied to all ambiguous HGR trials. Ambiguous HGR SVMs were evaluated with leave-one-out cross-validation, so that all scores were obtained from held-out trials. Each classifier returned a decision value (score) per ambiguous test trial, defined as the signed projection of the trial’s feature vector onto the hyperplane, which indicates how strongly the trial belongs to a class. Scores were averaged across repetitions and partitions, and the predicted label was the sign of the averaged score. Each ambiguous trial thus had three categorizations: the label predicted by the extreme HGR SVM (further referred to as *neural percept*), the label predicted by the ambiguous HGR SVM, and the *behavioral percep*t (the category identified by the participant). We compared these categorizations using the area under the ROC curve (AUC), which was computed against the behavioral percept by sweeping a threshold across the pooled SVM scores which were z-scored per channel. Because AUC is independent of the decision threshold, it allows comparison of the predictions of the ambiguous and extreme HGR SVMs.

Discrimination performance of the pooled AUC of ambiguous HGR SVMs was tested against a null distribution from 10,000 permutations of the behavioral percept labels, shuffled within participant, with the same permutation applied to all channels of a participant. Results showed that the behavioral percept could be predicted above chance (AUC = 0.529, p = 0.0145). We computed the difference between the AUCs of the extreme HGR SVMs and the ambiguous HGR SVMs per channel and bootstrapped from 10,000 resamples to obtain a 95% confidence interval (CI). The AUC tended to differ slightly between the ambiguous and extreme HGR SVMs (ΔAUC = 0.020; CI=[−0.008, 0.047]; Wilcoxon p = 0.0836). To compare the discrimination axes of the two SVM types, we computed the angle θ (0-180°) from the cosine similarity between the weight vectors averaged across partitions and repetitions. We tested whether the mean angle was equivalent to 90° using two one-sided tests (TOST equivalence test; also see^69^) against *a priori* margin of ±10°, implemented as a percentile. Equivalence was concluded if the 90% CI of the mean deviation from 90° fell entirely within the margin (10,000 resamples bootstrap over channels) The mean angle between the discrimination axes of the two SVM types was equivalent to 90° within ±10° (mean θ = 90.4°; mean deviation = 0.4°, 90% CI [−4.3°, 5.1°]; TOST p < 0.001). These results may suggest that the sensory axis of the extreme HGR SVM (/da/ vs. /ga/ sound) and the perceptual axis of the ambiguous HGR SVM (perceptual interpretation of the ambiguous sound) encode different, complementary qualities of ambiguous sounds although this finding needs to be interpreted with caution and corroborated in future studies.

We then used the trained SVM to address our main question: does pre-stimulus phase influenced neural perception, that is, the classification of the ambiguous HGR as /da/ or /ga/? To this end, we used analyses analogous to those for phase-dependent behavioral perception (Figure S3). In the first-level logistic regression, the behavioral percept was replaced with the neural percept (i.e., classification labels of the extreme HGR SVM). As before, the independent variables were the sine and cosine of the pre-stimulus phase, the channel, the participant, and their interaction. To account for variability in the quality of the ambiguous representation that is lost by class binarization, we included the trial-specific absolute decision value of the ambiguous HGR SVM as a fixed factor. As before, we extracted expected χ² values from the channel-specific null distribution (1000 permutations) and applied a boxcox transform to obtain the normalized χ²* distribution. At the second level, we applied the same multilevel linear mixed-effects (LME) analysis as for behavioral phase-dependent percept. A likelihood ratio test was used to derive the second-level χ² statistic from the mean χ² values across all time–frequency points within the significant cluster of the phase-dependent behavioral percept analysis. Significance was tested against a null distribution (n = 10,000) generated by repeating the multi-level model with permutations of the χ²-type factor (observed, expected). To assess the similarity between behavioral and neural percepts, we averaged the first-level χ² values within the behavioral cluster per channel and min-max normalized them separately for the behavioral and neural percept. We tested their association across channels with Spearman’s rank correlation.

#### Phase amplitude coupling (PAC)

From the pre-stimulus data, we took the phase of the low-frequency cortical response (2-30 Hz), and the power of the high-frequency cortical response (30-160 Hz). As PAC measure we used the modulation index (MI), a measure based on investigating a shift from a uniform distribution based on binned data ^36^. Data was binned in 20 equally spaced phase bins and for each bin we estimated the mean high frequency power. The logic of the analysis was the same as for the *phase-dependent categorization* analysis (see Figure S3). As a first level analysis, we created null-distributions for further testing by permuting the trial label of which phase of the low-frequency phase estimates (across the whole pre-stimulus window from -1 to 0 s) belonged to which trial per channel. We applied a boxcox transform generating observed MI* and expected MI* (mean of the transformed data). Then, the same lme model as in the *phase-dependent categorization* analysis was performed (including the boxcox transformation and multiple comparison correction) but using the MI*-type (observed vs expected) as independent variable. As PAC was overall strong for our sound-responsive channels, we also performed single channel statistics in which we used cluster-based correction over time-frequency points using the MI* as input.

#### Gamma clustering phase – da/ga tuning analysis

We related the theta phase at which gamma clustered in the pre-stimulus period to the /da/-/ga/ tuning of individual channels. To quantify gamma phase clustering, high gamma power (80-160 Hz) was averaged within 20 equally spaced theta phase bins (for phases estimated between 2 and 8 Hz). The gamma clustering phase of each channel was defined as the bin with maximal high-gamma power. To estimate da/ga tuning we extracted the mean HGR difference between the /da/ and /ga/ extremes in an early post-stimulus window (0.05-0.25 s post-stimulus). The relationship between gamma clustering phase and tuning we performed a lme model similar as in the *Phase-dependent categorization.* In this model, the dependent variable was the HGR difference, the fixed factors were the sine and cosine of the gamma clustering phase, and participant, channel and their interaction were included as random factors. Finally, we repeated this analysis twice, but using the tuning to the perceived categories of ambiguous sounds instead of the tuning to extreme sounds as dependent variable. In both instances, tuning was defined as the difference in mean HGR (0.05-0.25 s) between trials perceived as /da/ versus /ga/, but the grouping of the trials differed per analysis. In the first one, the HGR difference was based on /da/ versus /ga trials as classified by the participant (behavioral percept). In the second analysis, the / da/ and /ga trials were defined by the classification of withheld data by the ambiguous-sound HGR SVM. For all analyses, we controlled for multiple comparisons across theta frequencies using an FDR correction.

## Data Availability

According to patient’s clinical data privacy conditions, deidentified copies of the raw data supporting the conclusions presented in this study can be provided upon reasonable request. Please contact the corresponding author via email with any inquiries about the data.

## Author contributions

Conceptualization: SO. Methodology: SO. Formal analysis: SO, JP. Investigation: SO, JR, MEA-M, MR, JP. Resources: EG, RR, VK-M. Data Curation: SO, JR, MEA-M, EG, RR, VK-M, MR, JP. Writing – Original draft: SO, JP. Writing – Review & Editing: all authors read the last version. Project administration: SO, JR, MEA-M, EG, RR, VK-M, MR, JP. Funding acquisition: SO, RR, VK-M.

