## Supplementary Materials for "Item-specific clustering of high gamma-band activity on theta phase during speech categorization"

### Supplementary Information

#### Supplementary figures and tables

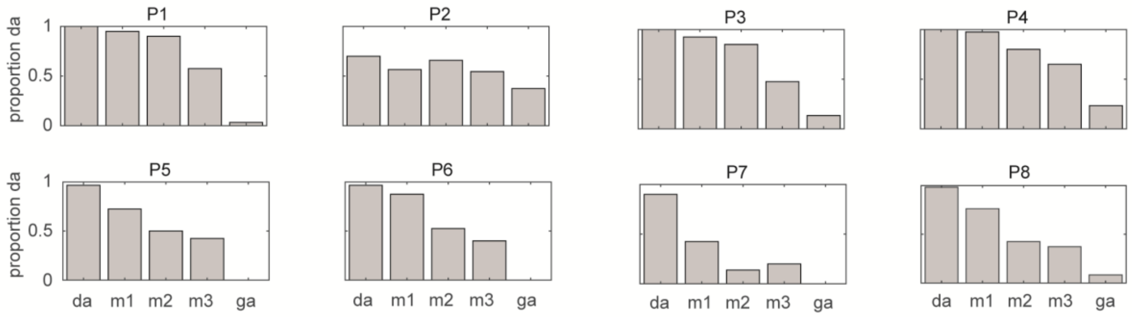


*Supplementary Figure S1*. Proportion /da/ responses for all eight participants for all morphs used. Note that participants 1 and 2 had slightly different morphs compared to the other participants.

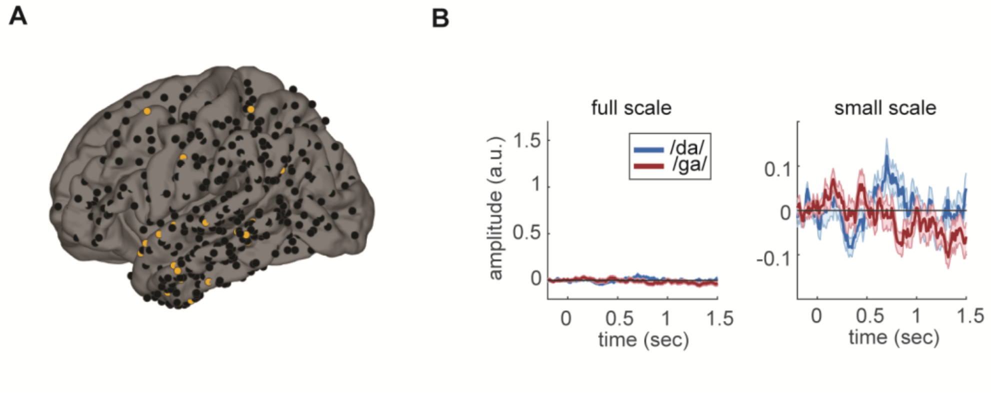


*Supplementary Figure S2*. Least response channels. A) The distribution of the channels with the weakest absolute HGR stimulus (0 to 1 s post-stimulus onset). B) Baseline-corrected high gamma response for the channels in A on the same scale as in the main manuscript (left) and on a small scale (right). Error bars reflect the standard error of the mean.

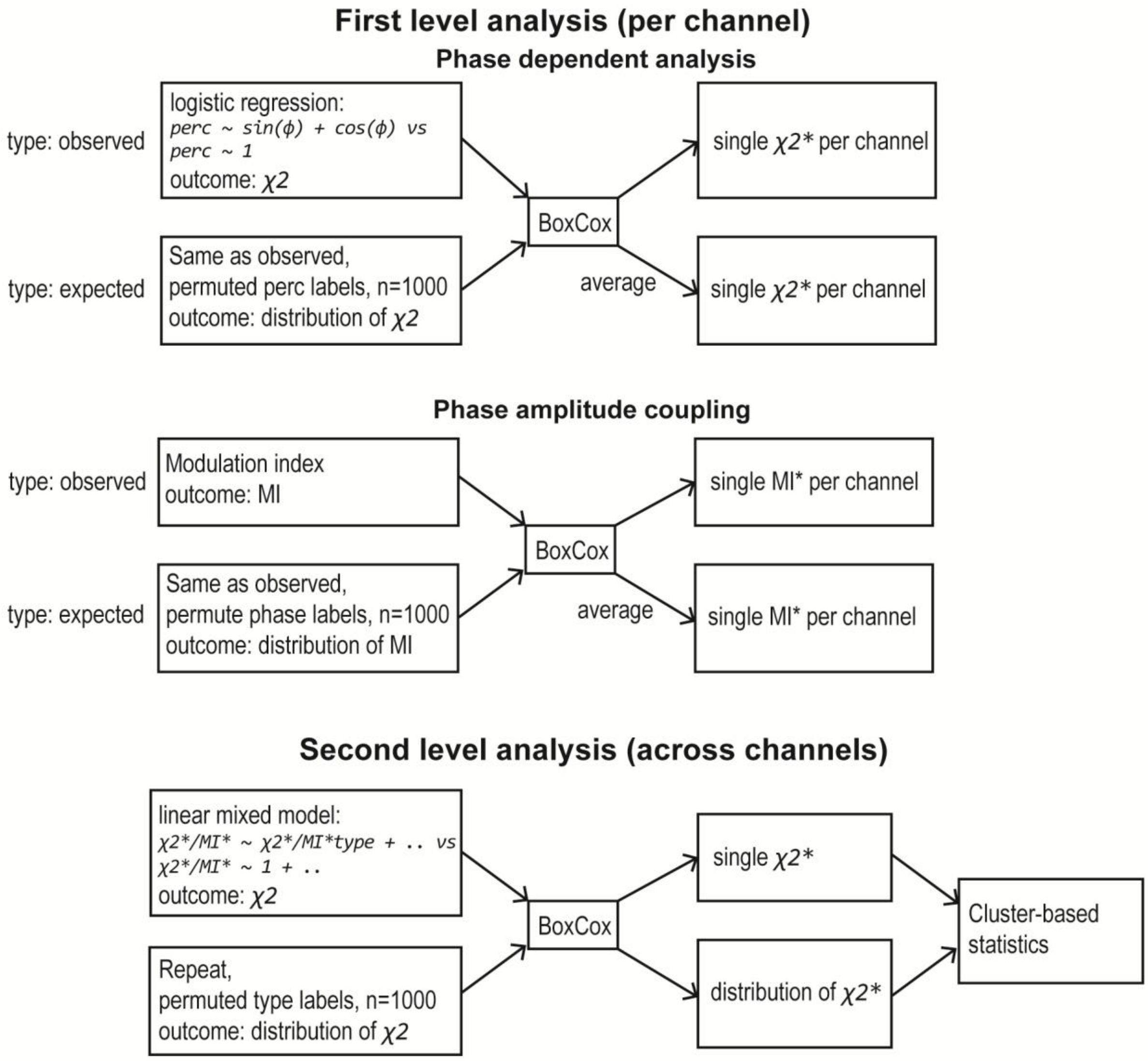


*Supplementary Figure S3*. Logic of the phase dependent analysis and the phase amplitude coupling analysis. On the first level analyses an observed and expected value is calculated either by regression or calculating the modulation index (MI) per channel. The phase dependent analysis for the behavioral and neural percept is identical except for an added independent variable (prediction score for ambiguous HGR SVM) in the neural percept logistic regression (see methods). As the $\chi^2$ and MI can be skewed, we applied a BoxCox transform. The first level analysis provides for each channel an observed and expected $\chi^2$*/MI*. On the second level analysis we apply a linear mixed model (lme) across with $\chi^2$/F/MI type (observed/expected) as factor. Random factors are not visualized here. Again, we performed permutations and a BoxCox transform and subsequent cluster-based statistics.

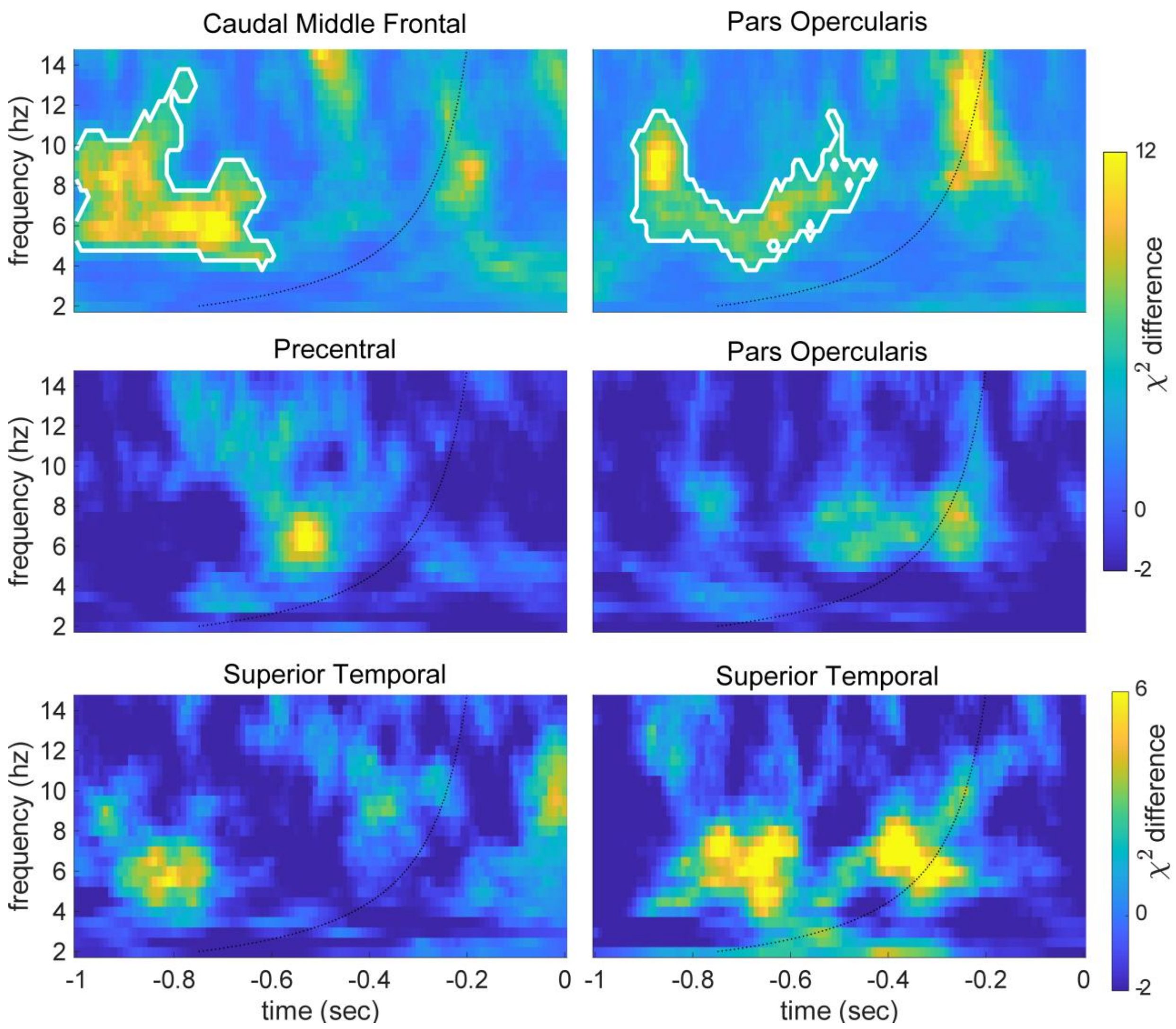


*Supplementary Figure S4.* Pre-stimulus phase determines the neural percept of an ambiguous sound. The outcome of the observed minus the expected $\chi^2$ from the logistic regression for a second set of example channels in various brain regions. Clusters with white outline: ($p < 0.05$, of the figure). Layout identical to Figure 2B.

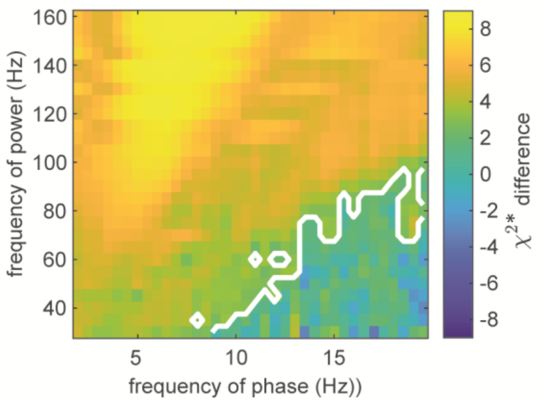


*Supplementary Figure S5.* Phase amplitude coupling result of the linear-mixed effect model. The main cluster ($p < 0.001$, white outlines of the figure) of the phase-amplitude coupling analysis encompasses a large section with strongest difference for theta and high gamma power.

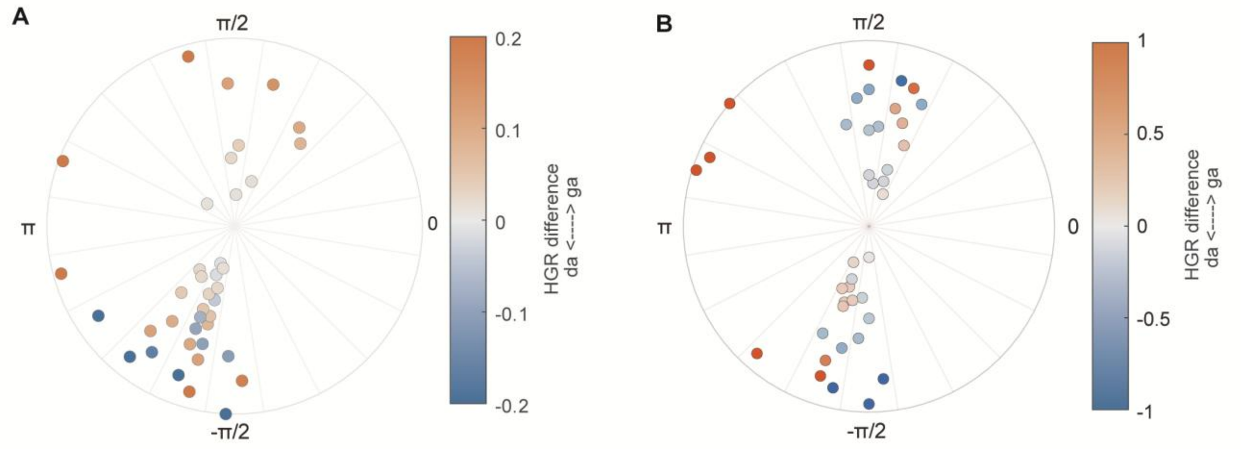


*Supplementary Figure S6.* Additional gamma clustering-tuning relationship. A) Rose plot of gamma response difference to the extreme sounds for 4.5 Hz. B) Rose plot for the difference in the SVM class predictions at 6.5 Hz. Conventions are the same as in Figure 4.

**Supplementary Table 1**

Significant high gamma response channel labels

| Label | Hemisphere | Count |
|---|---|---|
| Superior Temporal | Left | 12 |
| Supramarginal | Left | 4 |
| Pars Opercularis | Left | 4 |
| Inferior Temporal | Right | 3 |
| Inferior Temporal | Left | 2 |
| Postcentral | Left | 2 |
| Precentral | Left | 2 |
| Rostral Middle Frontal | Left | 2 |
| Bankssts | Left | 1 |
| Caudal Middle Frontal | Left | 1 |
| Middle Temporal | Left | 1 |
| Superior Parietal | Left | 1 |
| Postcentral | Right | 1 |
| Precentral | Right | 1 |
| Precuneus | Right | 1 |
| Superior Temporal | Right | 1 |
